# *R3DFEM*: an R package for running the 3D-CMCC-FEM model

**DOI:** 10.1101/2024.10.07.616968

**Authors:** Elia Vangi, Daniela Dalmonech, Alessio Collalti

## Abstract

Forest ecosystems account for about one-third of the Earth’s land area, and monitoring their structure and dynamics is essential for understanding the land’s carbon cycle and their role in the greenhouse gas balance. In this framework, process-based forest models (PBFMs) allow studying, monitoring and predicting forest growth and dynamics, capturing spatial and temporal patterns of carbon fluxes and stocks. The ‘Three Dimensional-Coupled Model Carbon Cycle - Forest Ecosystem Module’ (3D-CMCC-FEM) is a well-known eco-physiological, biogeochemical, biophysical process-based model, able to simulate energy, carbon, water and nitrogen fluxes and their allocation in homogeneous and heterogenous forest ecosystem. The model is specifically designed to represent forest stands, from simple ones to those with complex structures, involving several cohorts competing for light and other resources in a prognostic way. Currently, the model is implemented in C-language, which can be challenging for the broad public to use, and thus limiting its applications. In this paper, we present the open-source R package ‘R3DFEM’ which introduces efficient methods for: i) generating and handling input data needed for the model initialization; ii) running model simulations with different set up and exploring input; and iii) plotting output data. The functions in the R-package are designed to be user-friendly and intended for all R users with little to advanced coding skills, who aim to perform simulations using the 3D-CMCC-FEM. Here we present the package and its functionalities using some real case study and model applications.

## 1. Introduction

Forest ecosystems play an active role in the global carbon cycle, acting as a climate regulator by modulating the exchange of energy, carbon and water fluxes between lands and atmosphere (Huntingford et al., 2009; Collalti et al., 2016). In particular, via the gross primary production (GPP), forests fix atmospheric CO_2_ as organic compound offsetting anthropogenic emission of greenhouse gases. Due to the key role of forests in the climate change context, much progress was achieved in the development of process-based forest models (PBFMs) integrating more and more representation of detailed eco-physiological and population-related processes (Mäkelä et al., 2000). However, most models have limitations in accurately predicting forest photosynthesis, growth and carbon dynamics, particularly for forests that exhibit high structural complexity Natural or semi-natural forests, especially in the Mediterranean regions, can be composed of numerous tree species with complex horizontal and vertical structures, resulting from past management and disturbance regime, which, in turn, causes complex interactions among trees and different light conditions. Despite the importance of representing forest complexity, just a few models are able to represent heterogeneous ecosystems (Seidl et al., 2012; Collalti et al., 2014; De Wergifosse et al., 2022), such as those in the Mediterranean areas. The 3D-CMCC-FEM model is specifically designed to represent forest stands of different dimensions (from the common hectare to the 1 km x 1 km scale) and from simple ones to those with complex structures, involving several cohorts competing for light and other resources in a prognostic way, that is, not constrained by diagnostic observations over time but rather builds over a series of first principles and initial data representing the initial conditions of the system. The model is developed with the aim to simulate both fluxes and stocks and is able to capture dynamics occurring in homogeneous and heterogeneous forests with different tree species, for different ages, stem diameters and tree height classes competing for resources (e.g. light, water). The 3D-CMCC-FEM simulates carbon fluxes, in terms of gross and net primary production (GPP and NPP, respectively), partitioning and allocation in the main plant compartments (stem, branch, leaf, fruit, fine and coarse root, including non-structural carbon compounds i.e. a reserve carbon pool (Merganicova et al., 2019; Collalti et al., 2019). In the latest versions (see for example: Collalti et al., 2019, 2020; Dalmonech et al., 2022, 2024; Testolin et al., 2023; Morichetti et al., 2024; Vangi et al., 2024a and 2024b), nitrogen fluxes and allocation, in the same carbon pools, are also considered. The 3D-CMCC-FEM as a stand-level model, is initialized providing the forest structure information, such as species share, average diameter at breast height (DBH) and age class. In turn, initializing with current observed forest structure, the model implicitly embeds the effect of past management practices and disturbance, overcoming the need to know the exact history of the site in the classical spin-up and transient simulation approach, used in e.g. global or regional vegetation models. Nevertheless, the 3D-CMCC-FEM model also implements past management practices (e.g. thinning and harvest) and can predict their effects on forest growth and carbon sequestration and stock under future climate change scenarios, or within a ‘what if’ scenarios framework.

The 3D-CMCC-FE is written in C-language and divided into several libraries and source files, each describing the main physiological processes, within thousands of line codes. Despite the documented potential of the model in monitoring and forecasting forest ecosystems in simulating forest growth under different management and climate assumptions, its applicability still remains for users with a good level of programming, limiting the possibilities offered by this tool. Wrapping the model in an R package can ensure a simpler approach for users with less programming experience and a better way to share the knowledge on which the model is based, expanding the user base and simultaneously improving the model itself through the reporting of issues and the experiences of researchers.

This paper aims to: i) present the *R3DFEM* R package for running the 3D-CMCC-FEM model; and, ii) test the package in different real-case scenarios. First, we provide a detailed description of the functions and features (section 2). Second, we show illustrative examples by applying *R3DFEM* over different forest stands and validate the outputs against field measurement (section 3). Third, we explore the impact of this new package by highlighting the scientific and operative contribution of *R3DFEM* (section 4).

## 2. Design and implementation

*R3DFEM* is an R package written in R 4.2 (R Core team 2017) designed as a wrapper to the main source C-code of the 3D-CMCC-FEM model. Currently, the main source code is compiled in an executable (.exe) file for Windows OS only and the package, despite being tested also for iOS and UNIX, the *R3DFEM* is guaranteed to be compatible only for Windows. At the installation of the package, all the routines and process, written in C-language in separate files are downloaded. The user does not need to interact with these files, since they are all already compiled into the .exe file. The R package follows a simple name convention: all function names start with a verb indicating the function’s primary purpose followed by an underscore (i.e. plot_, run_, check_). *R3DFEM* uses the *data.table* package (https://r-datatable.com) for the data structure, allowing fast and memory-efficient data aggregation and manipulation. The main function (run_3DFEM) is designed to check the input data and call the .exe in a way that is easily parallelizable exploiting the computational power of the modern PC. The output of the 3D-CMCC-FEM is a simple txt file that, in addition to the in-built functions in the package, can be read by the most common R packages for data handling and plotting, such as *data.table*, *readxl* and *ggplot2*. *R3DFEM* provides functions for: i) checking and creating input data, ii) running simulations, iii) plotting input and output data. For each function we provide detailed information in subsequent sections.

### 2.1. Inputs and outputs

Below we present a schematic description of the input needed by the functions in the R package:

For initialization, the 3D-CMCC-FEM requires as input data:

- The initial stand conditions: species name (since the model is parameterized at specie-level), age, mean tree height, diameter at breast height (DBH), number of trees per size cell. The initial data are aggregated per classes (height classes, cohorts and species) by a pre-processing activity as following: (1) the relative values of diameters class is associated for each species, (2) the corresponding value of height class is assigned for each diameter class, and (3) the relative age is assigned for each height class (Collalti et al., 2014; 2024).
- Species-specific parameters, which are mostly based on species-specific eco-physiological and allometric characteristics and can be partially derived from forest inventories and literature (Collalti et al., 2019). Along with the package comes a suite of already parameterized files for different and most common European tree species, used in many real case studies across Europe.
- Meteorological forcing data: daily maximum (Tmax, °C) and minimum air temperature (Tmin, °C), soil temperature (Tsoil, °C), vapour pressure deficit (hPa), global solar radiation (MJ m^−2^ day^−1^) and precipitation amount (mm day^−1^).
- Annual atmospheric CO_2_ concentration and nitrogen deposition (optional)(Collalti et al., 2018).
- Soil and topographic information: soil depth, average sand, clay, silt percentages and elevation.

All input data need to be written into separate .txt files whose structure is fully described in the user manual (https://www.forest-modelling-lab.com/_files/ugd/8a7700_d31451e9a5e64073b50c07f7f007eb71.pdf). Based on the input files and the argument setting, the function wrapping the model (run_3DFEM) creates a setting file in the output directory which is used only by the internal C code; the user does not need to interact with the setting file.

The main output of the 3D-CMCC-FEM (either at daily, monthly or annual scale) are: Gross Primary Productivity (GPP), Net Primary Productivity (NPP), and state variables such as evapotranspiration (ET), Leaf Area Index (LAI) and rain interception (to cite some). Results are obtained either at class-level (species, diameter, height, or age class level), layer-level (as sum of all tree height classes in the same layer), and grid level (as sum of all classes in the different layers). The model provides information to support decision-making in forest management planning, such as mean annual volume increment (MAI), current volume increment (CAI), basal area, and DBH.

Since this paper has not the aim of describing the model processes and functionalities, for detailed information about 3D-CMCC-FEM and its applications we strongly encourage to refer to the literature (Collalti et al., 2014, 2016, 2020, 2024; Marconi et al., 2017; Mahnken et al., 2022; Dalmonech et al., 2022, 2024; Testolin et al., 2023; Vangi et al., 2024a, 2024b; Morichetti et al., 2024), and the main web page (https://www.forest-modelling-lab.com/the-3d-cmcc-model, accessed online on 26/09/2024), where the most updated user guide can be found (which include the detailed description of all inputs and outputs, as well as the instruction for launching the model from command line, Eclipse and Bash; Collalti et al., 2022). Throughout the paper and in the description of the functions we will often refer to the user guide and the official web page.

### 2.2. Main functions

Below is shown a schematic representation of the main function of the package and their relations in respect to outputs and inputs (Figure 1).

**Figure 1.**
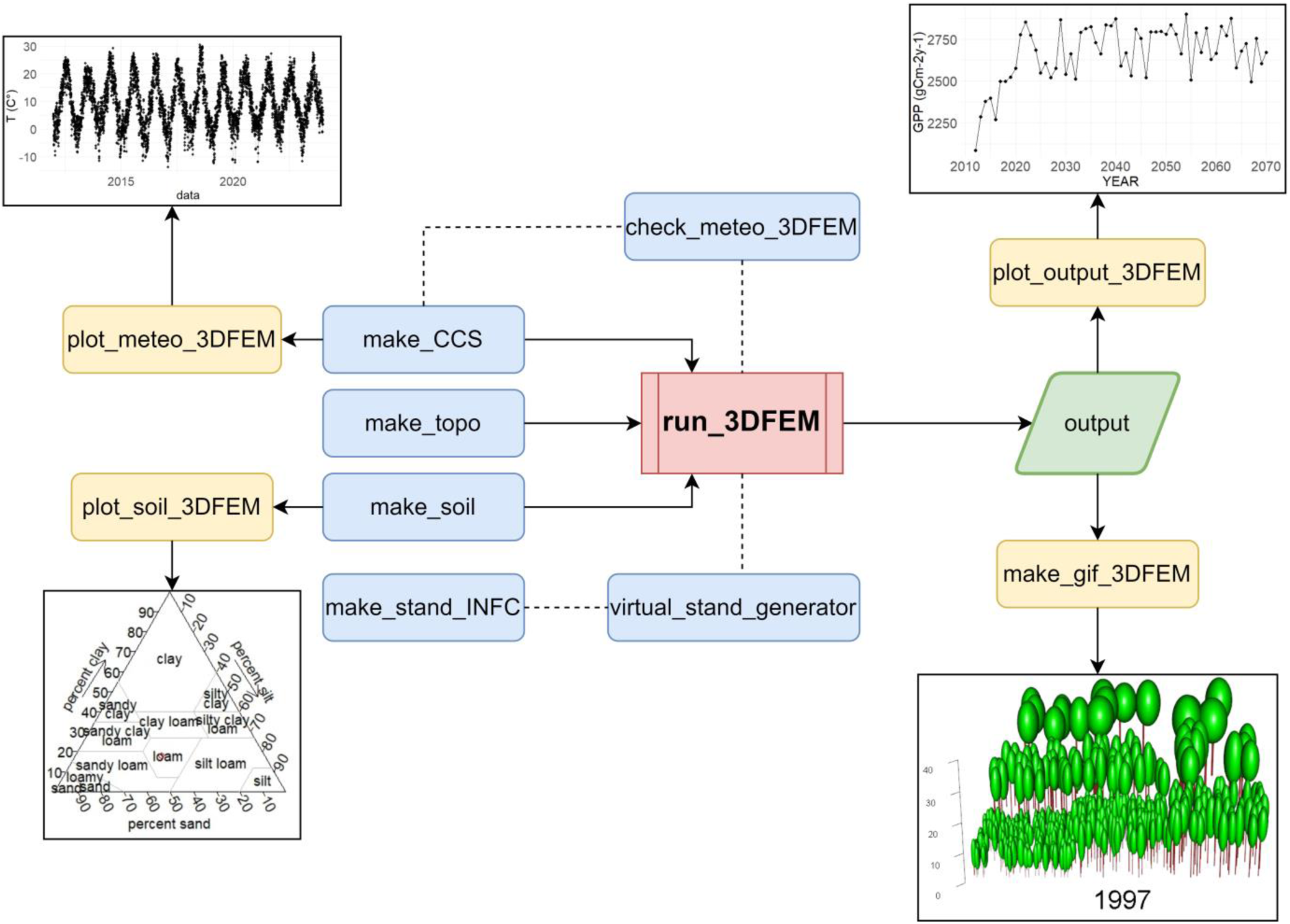
Flowchart of *R3DFEM* package. In blue, functions for creating/manipulating input data; in red, the main function for running the model; in green, the output; in yellow, functions for plotting inputs and outputs.

#### 2.2.1 initialization and input check

*R3DFEM* offers functions to check the requirements of input data for the model and some facilities to perform common tasks in modeling applications, such as detrending climate data for creating baseline scenarios in climate change impact studies.

The function *check_meteo_3DFEM* takes as input a path to a .txt meteo file (see the 3D-CMCC-FEM User Guide parag. 4.5 for the specific format and name convention of the meteo file) and checks several assumptions to ensure that the input meets the model requirements, such as the consistency of temperature values (Tmin<Tmean<Tmax), the consistency of solar radiation, precipitation and relative humidity values (values >0), the correctness of the number of days in each month and years (leap years are also considered), the presence of missing values or wrong column names spelling. The function reports any inconsistency with the model specification and (if specified by the user) tries to fix any errors.

The function *make_stand_INFC* uses the package in-built tree-level data from the Italian National Forest Inventory (INFC) to create the stand initialization file, meeting the model specification (see the User Guide parag. 4.2 for the format of the stand file). Like all the other input data, this is a txt file with the stand structural characteristics, such as the age, mean DBH, height, number of trees for the main species in the plot.

The function *virtual_stand_generator* creates “virtual stands” from a real stand file (such as one created by *make_stand_INFC*) by running the model and extracting the structural attribute from the simulation at specific ages of the stand. This function is useful to assess (and to depict) the effect of the age at the beginning of a simulation, for the same stand (see the “Composite Forest Matrix” description in Dalmonech et al., 2022) or create representative forest parcels, without the need to perform new field campaign or inventory.

The function *make_topo* creates the topographic file following the model specification, starting from the coordinates and elevation of the site (see the User Guide parag. 4.4 for the format and name convention of the topographic file).

The function *make_CCS* creates a “current climate scenario” from a meteo file, by detrending and repeating cycles of observed meteo up to a user-defined time span. This function is useful for creating baselines scenario i.e. counterfactual scenario, against which climate change scenarios and 3D-CMCC-FEM model outputs can be compared (see for an example Collalti et al., 2018).

#### 2.2.2. Running simulations

The main function of the package is *run_3DCMCCFEM* which is a wrapper around the C code compiled in the exe file provided within the package. The function allows to run the model from the R environment. Each argument of the C functions is matched in the wrapper, so that the model can be launched with every possible setting. First, the function performs several checks needed to ensure the consistency of all arguments specified by the user, then check the consistency against the model specifications and finally builds the system call to run the model, translating the R-code to a Bash call. All the inputs needed for the simulation (see the 3D-CMCC-FEM User Guide parag. 4 for a detailed overview of each input file), must be in the same directory, whose path is an input for the function. The output is saved locally following a root path that depends on the simulation setting (i.e. temporal scale of the output, name of the simulated site, whether the simulation has been performed with fixed CO_2_ or with active management, etc.) and is managed internally by the C-code. The user needs to specify the working directory where to save the simulation outputs and the function create the tree path accordingly. The output files saved by the function consist of the main output file, which contain fluxes and stocks values for each time step (i.e. day, month, year), each species and layer (see the 3D-CMCC-FEM User Guide pararg. 4.10 for the detailed output list), a debug file, where, in case of failed simulation, all the errors of the run and the list of the input file used for the simulation are reported (useful for debugging and sharing).

#### 2.2.3 Plotting

The package implements some functions for a visual assessment of inputs and outputs, which can be used also for publications, reporting and other activities.

The function *plot_soil_3DFEM* creates a soil texture diagram (also known as triangle plot or ternary plot) from the input soil file of the model (see the 3D-CMCC-FEM User Guide parag. 4.3 for detailed information on the soil input file). In a ternary plot, 3D textural coordinates, which sum is constant, are projected in the 2D space, using simple trigonometry rules. Our Package exploits the *triax.plot* function from the *plotrix* R-package (Lemon, 2006) to build the soil diagrams.

The function *plot_meteo_3DFEM* creates a time-series plot of climate forcing variables starting from the path of a meteo file. It is possible to define the time span of the time-series, and to plot a moving average at a user-defined time window.

The function *plot_output_3DFEM* allows to plot one or more variables from the output file of a simulation run. The function can plot the time-series of one or more output variables or a scatterplot of two variables, depending on the number of inputs specified in the function. In particular, the function accepts two arguments, *x* and *y*. If *y* is not specified the resulting plot will be a time-series plot, a scatterplot otherwise.

For examples of graph produced by the plot functions see the next parag 3.

## 3. *R3DFEM* applications

The following section describes the use of the *R3DFEM* package in real-case applications aimed at illustrating the main functions capabilities. In particular, in this section we will see: i) a validation against Eddy tower fluxes (Pastorello et al., 2020) at the Sorø forest site (Denmark); ii) the creation of virtual forest stands at the Hyytiälä forest site (Finland); iii) a comparison of a climate change scenario against the baseline “current climate” scenario at the Bilỳ Křiž forest site (Czech Republic).

### 3.1 Study sites

The study was conducted in three even-aged, previously managed European forest stands: i) the Boreal Scots pine (*Pinus sylvestris* L.) forest of Hyytiälä, Finland (FI-Hyy); ii) the wet temperate continental Norway spruce (*Picea abies* (L.) H. Karst) forest of Bílý Krìz in the Czech Republic (CZ-BK1); and iii) the temperate oceanic European beech (*Fagus sylvatica* L.) forest of Sorø, Denmark (DK-Sor). The chosen sites have been selected due to their long monitoring history and the availability of a wide range of data sources for both carbon fluxes and biometric data for model evaluation provided in the PROFOUND database (Reyer et al.,2020), as well as bias-corrected climate scenarios for simulations under climate change scenarios as provided within the ISIMIP initiative (https://www.isimip.org/). For more details about these sites please refer to Mahnken et al. (2022), Vangi et al. (2024a, 2024b) and Morichetti et al. (2024).

### 3.2. Case 1: Validation of GPP at Sorø

In this real-case application is shown the validation of the main flux (GPP, gC m^−2^ day^−1^) at the daily temporal scale for the beech forest of Sorø against the flux data measured by the eddy covariance tower from the FLUXNET database installed at the same site /fluxnet.org/data/fluxnet2015-dataset/).

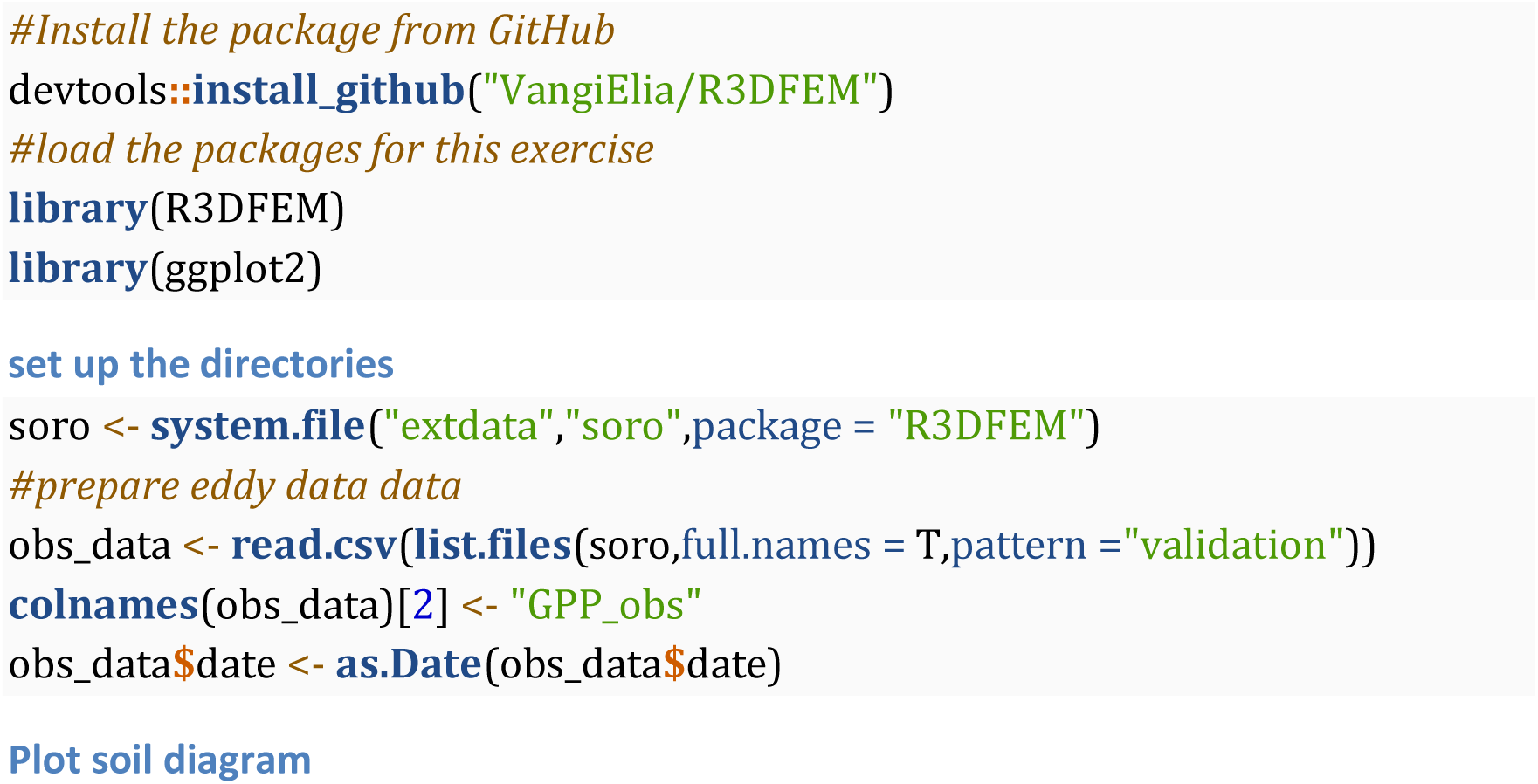

Here the function *plot_soil_3DFEM* is used to produce the soil diagram plot of the site, based on the input soil file (Figure 2).

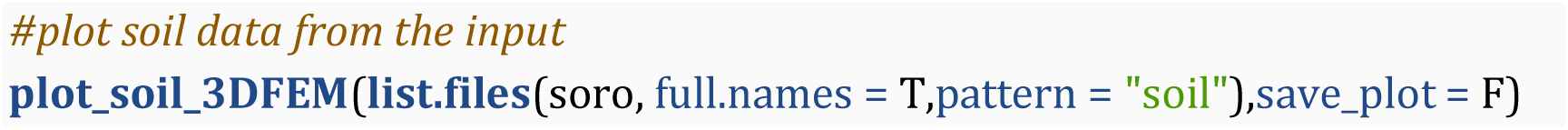

**Figure 2.**
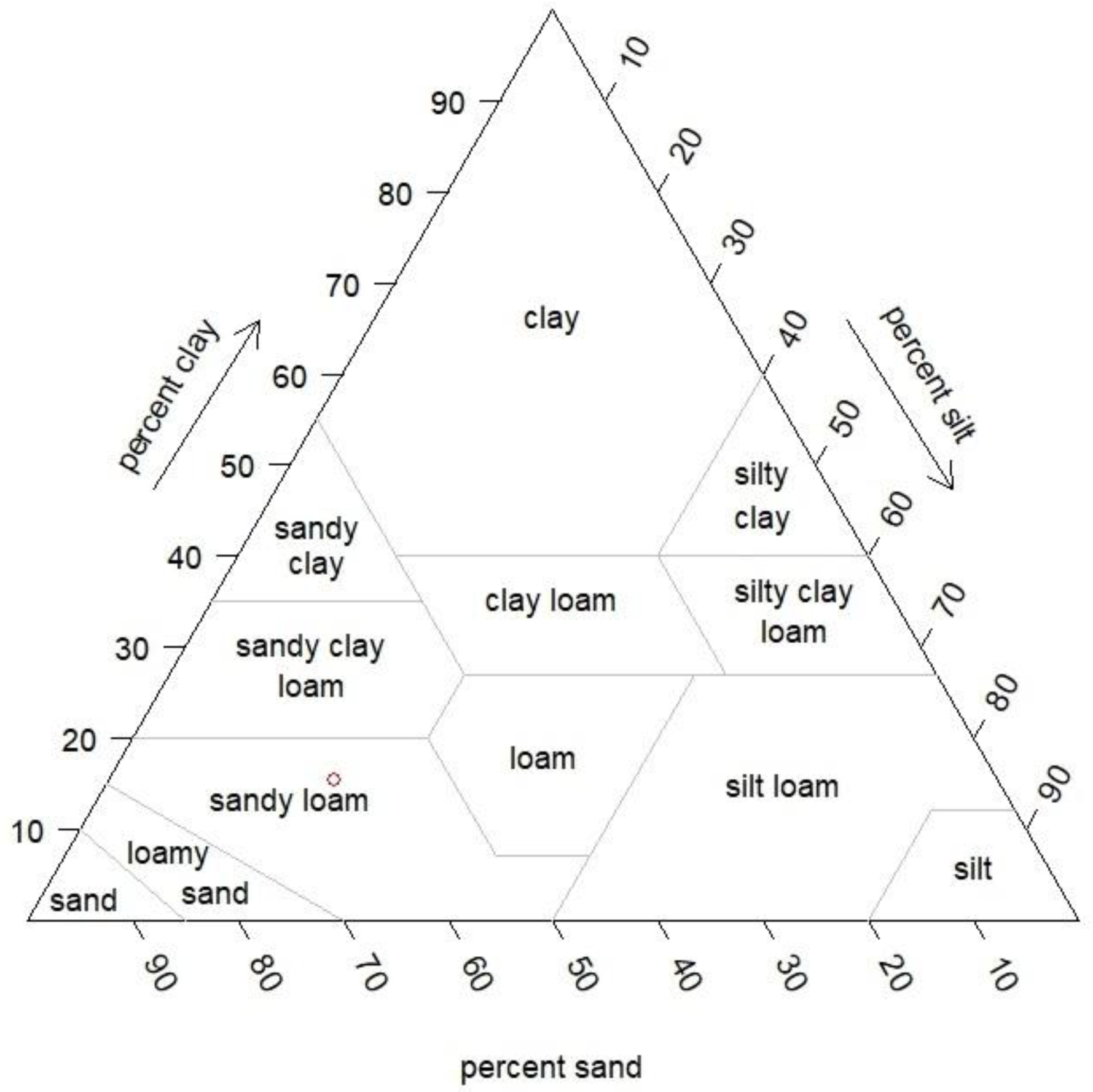
Soil texture diagram obtained with the function plot_soil_3DFEM at Sorø

### Run simulations

Here we run a simulation to compare the model output with the measured data from the eddy covariance tower. Since the measured data are provided for the period 1996-2014, and since the available climate data for the site start from 1997 the simulation period is 1997-2014, to cover as much as possible the time window of the observations.

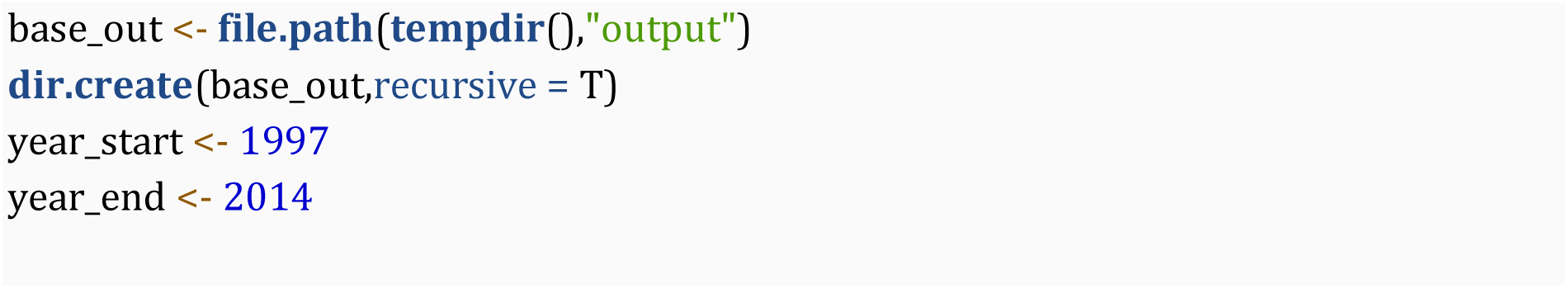
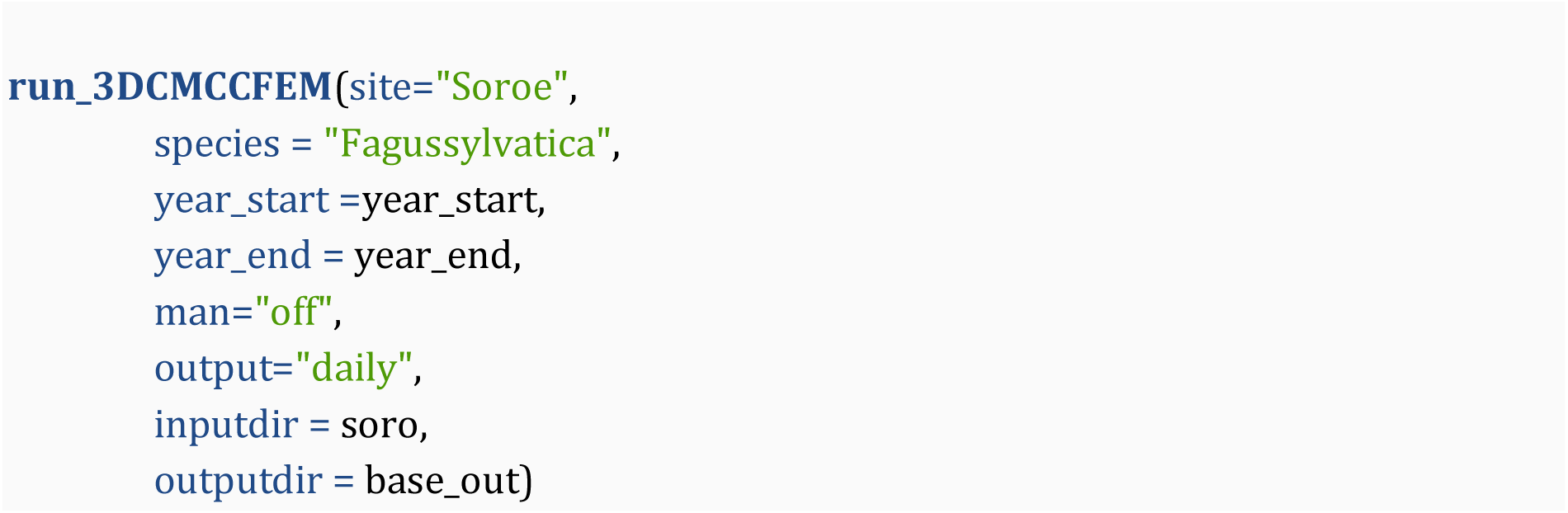

### Read and evaluate the output

In the next piece of code, the model output is read into the R-environment and the variable of interest is plotted against the measured data (Figure 3).

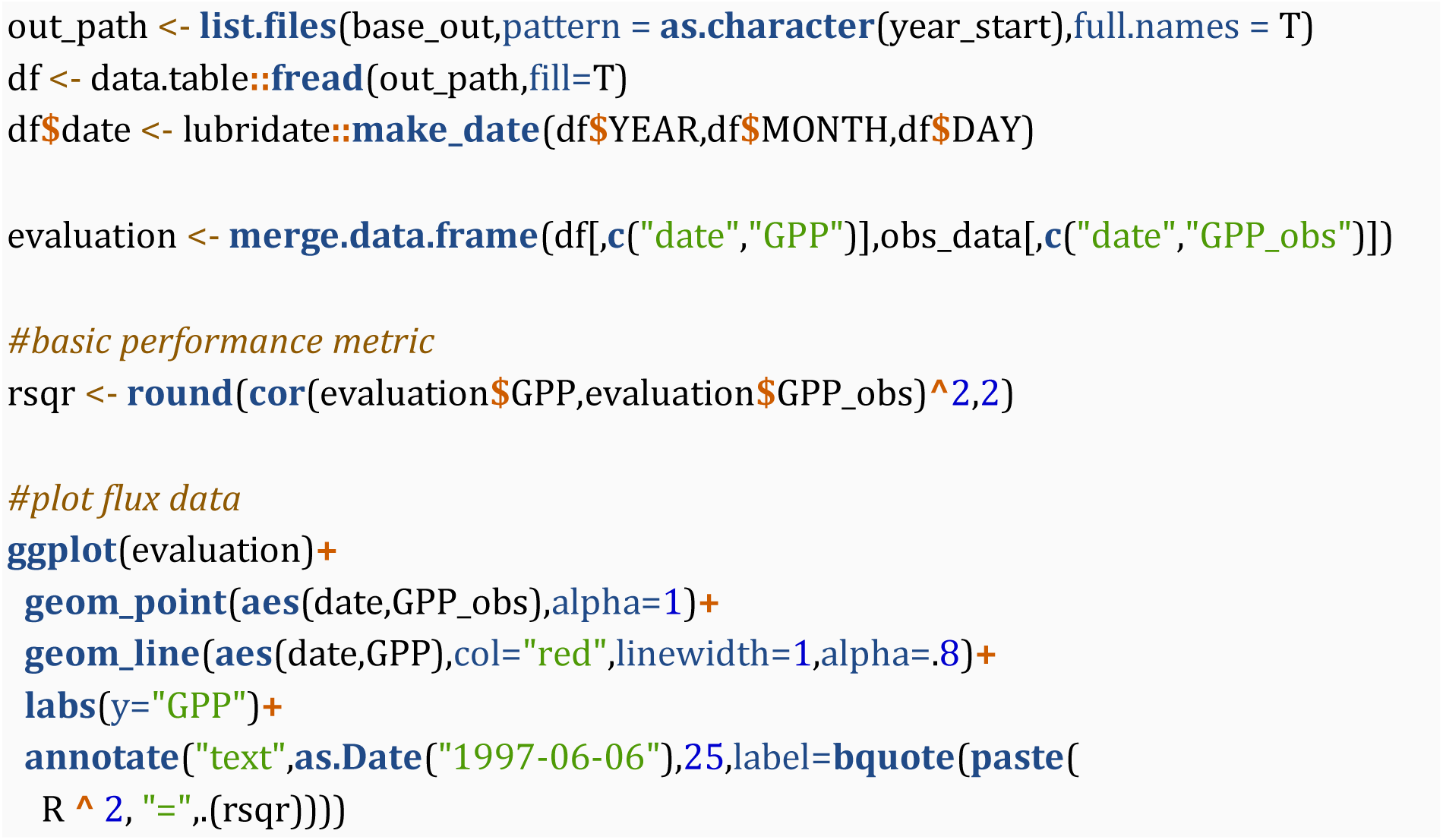

**Figure 3.**
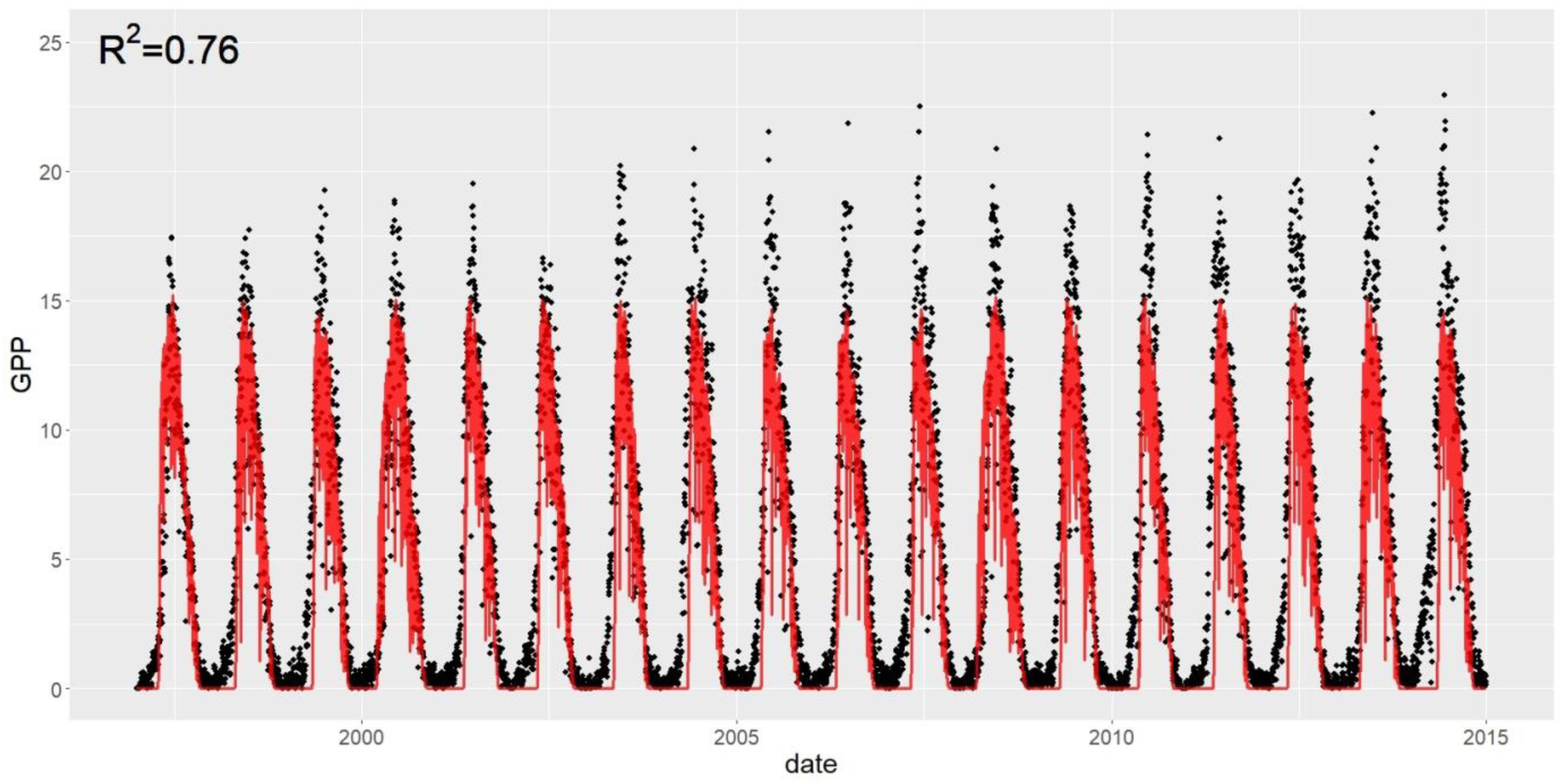
Comparison of daily GPP flux between model simulation (in red) and observed eddy covariance flux data (black dot).

### 3.3. Case 2: virtual stand generation at Hyytiälä

In this real-case application is shown the use of the *R3DFEM* package for the creation of virtual stands on the Hyytiälä site (Finland).

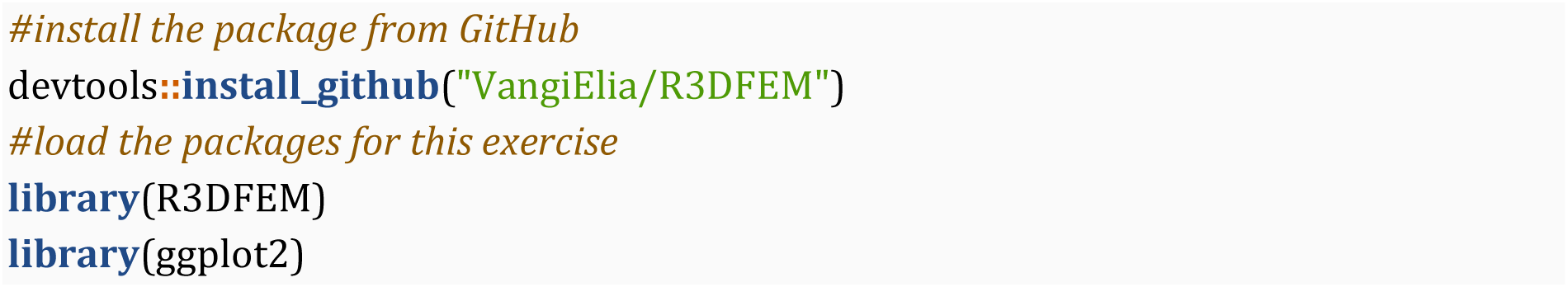

#### set up the directories

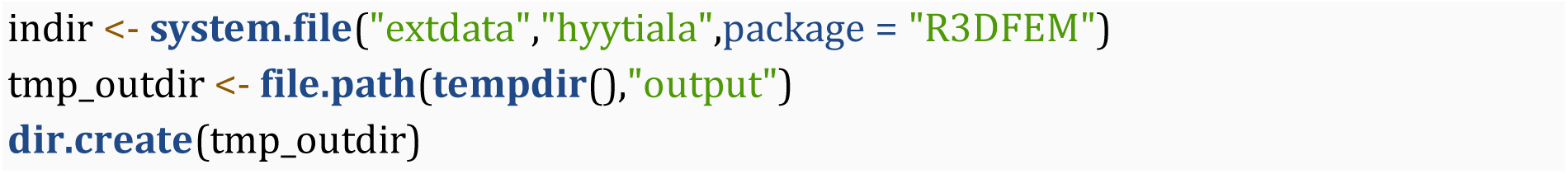

#### creates the “virtual stands”

The main function in this application is *virtual_stand_generator*. The function *virtual_stand_generator* creates a new folder in *outputdir* called “virtual stand” where the new virtual stands information are saved. In figure 4 are shown the structural variable of stand of different ages created with the *virtual_stand_generator* function.

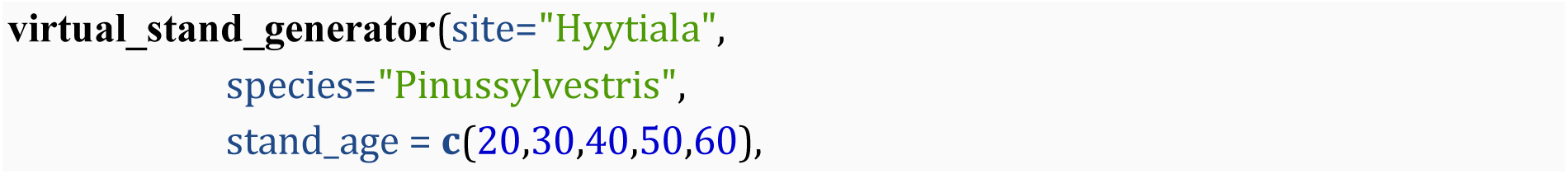
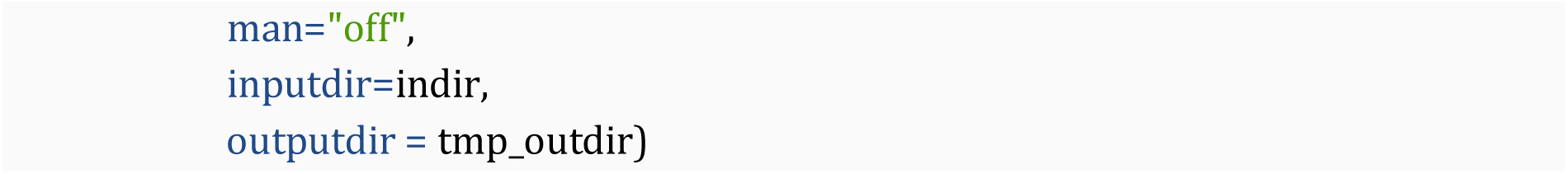

#### checks the output

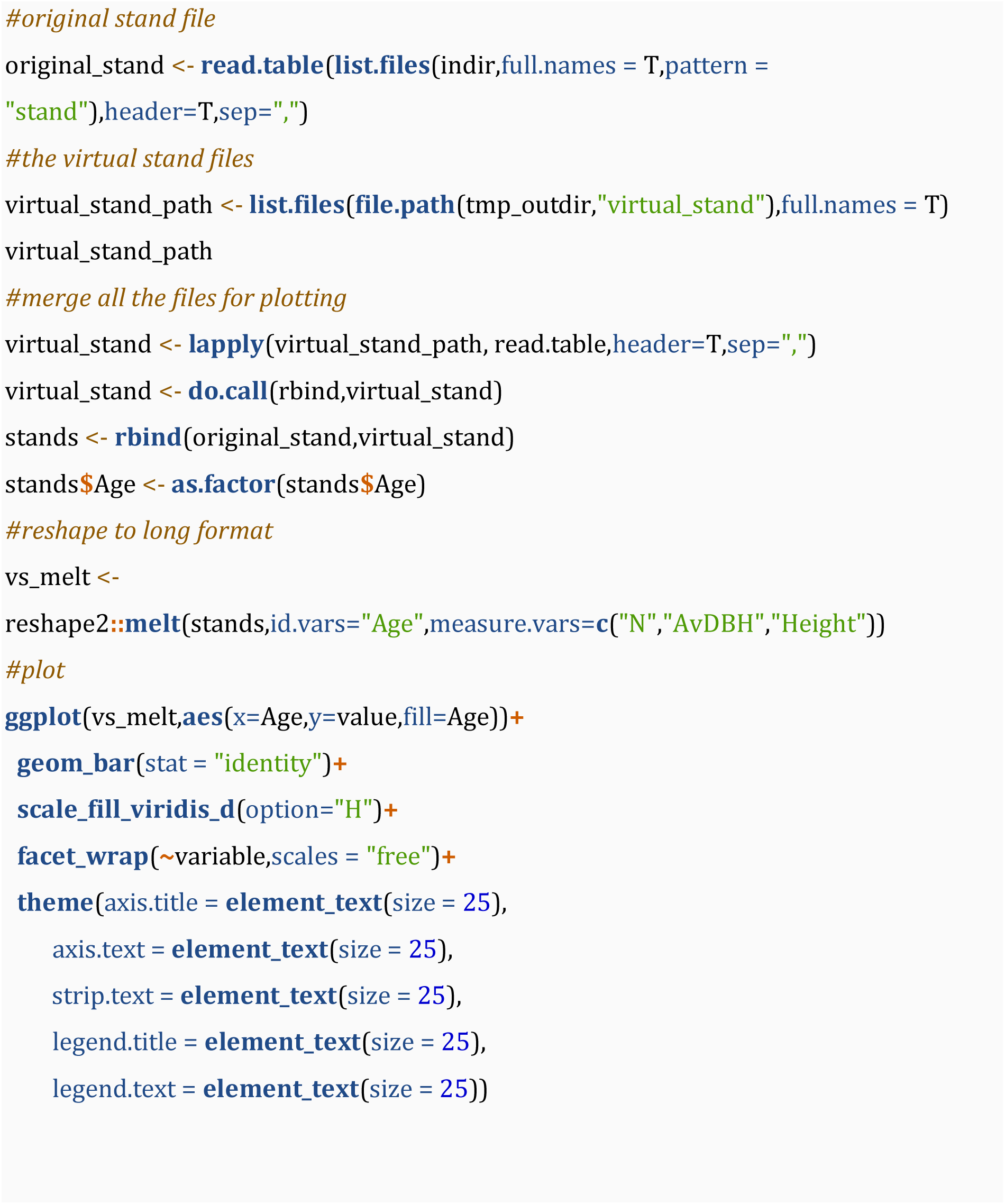

**Figure 4.**
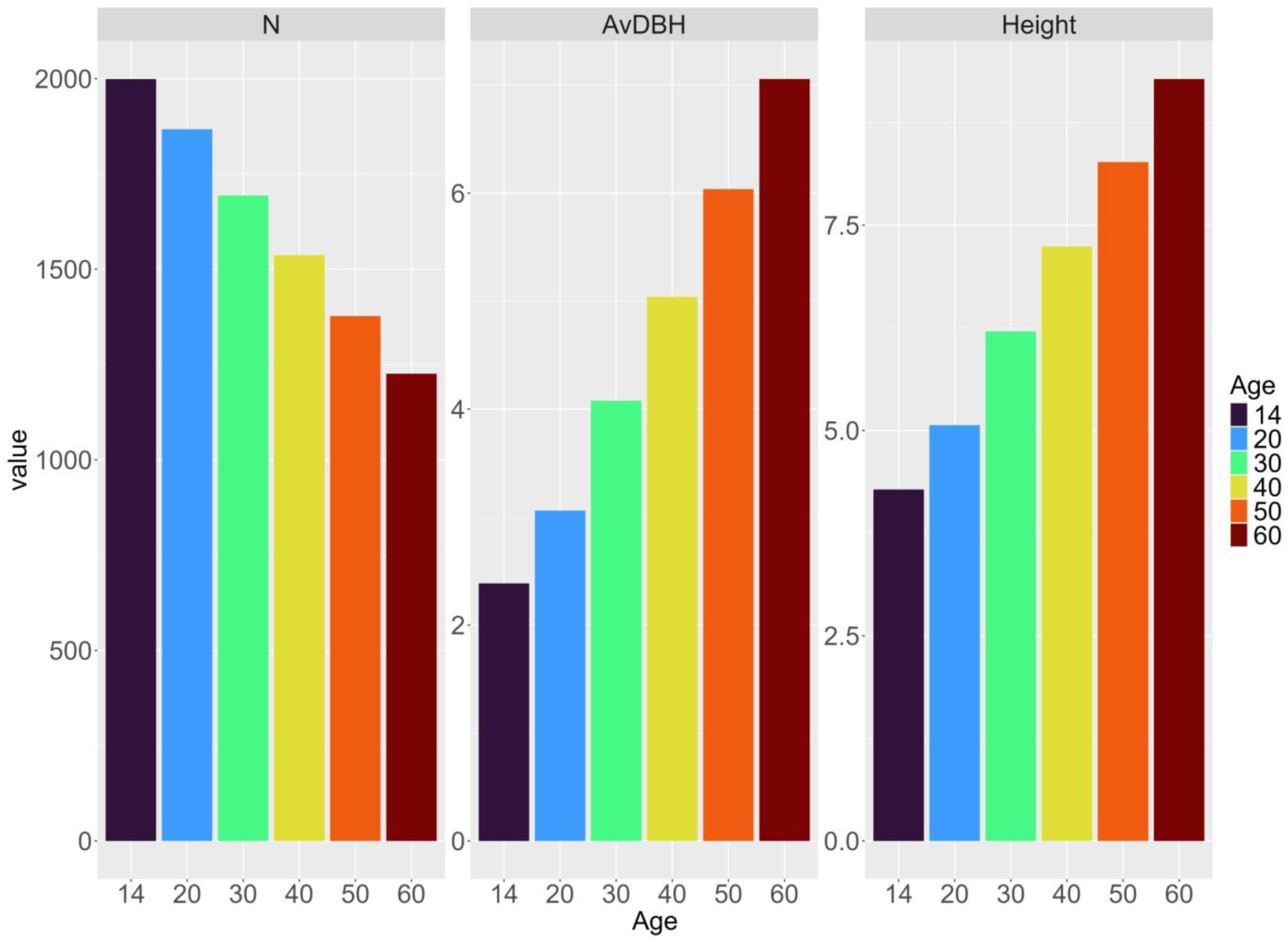
Structural variable of stand of different ages. The 14 years old stand is the original one from which the ”virtual stands” were created.

#### 3.4. Case 3: climate change scenario at Bilỳ Křiž

In this real-case application is shown the use of the *R3DFEM* package for running simulations under different climate change scenarios. We will use a baseline scenarios (created by de-trending and repeating observed climate for 100 years) and the RCP 8.5 scenarios, the most severe in terms of increase in temperature, solar radiation and atmospheric CO_2_ increase. The package is used also for plotting some input and model output data.

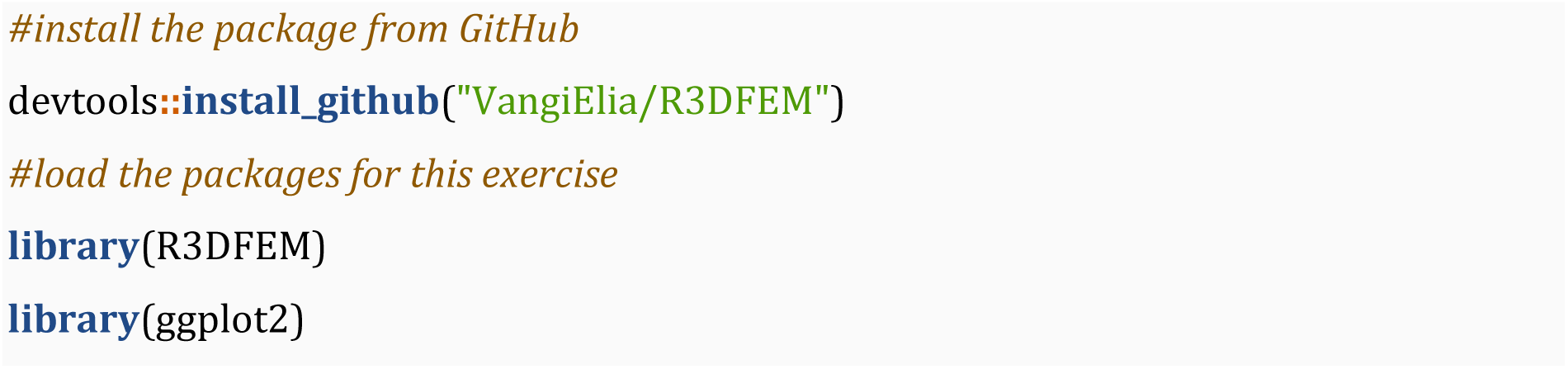

##### set up the directories

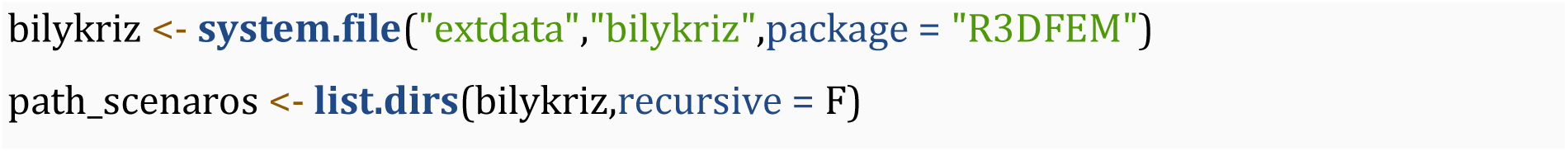

##### Plot mean temperature for both scenario

Here the function *plot_meteo_3DFEM* is used to produce the time-series plot of the mean temperature from the meteo file of both scenarios (Figure 5).

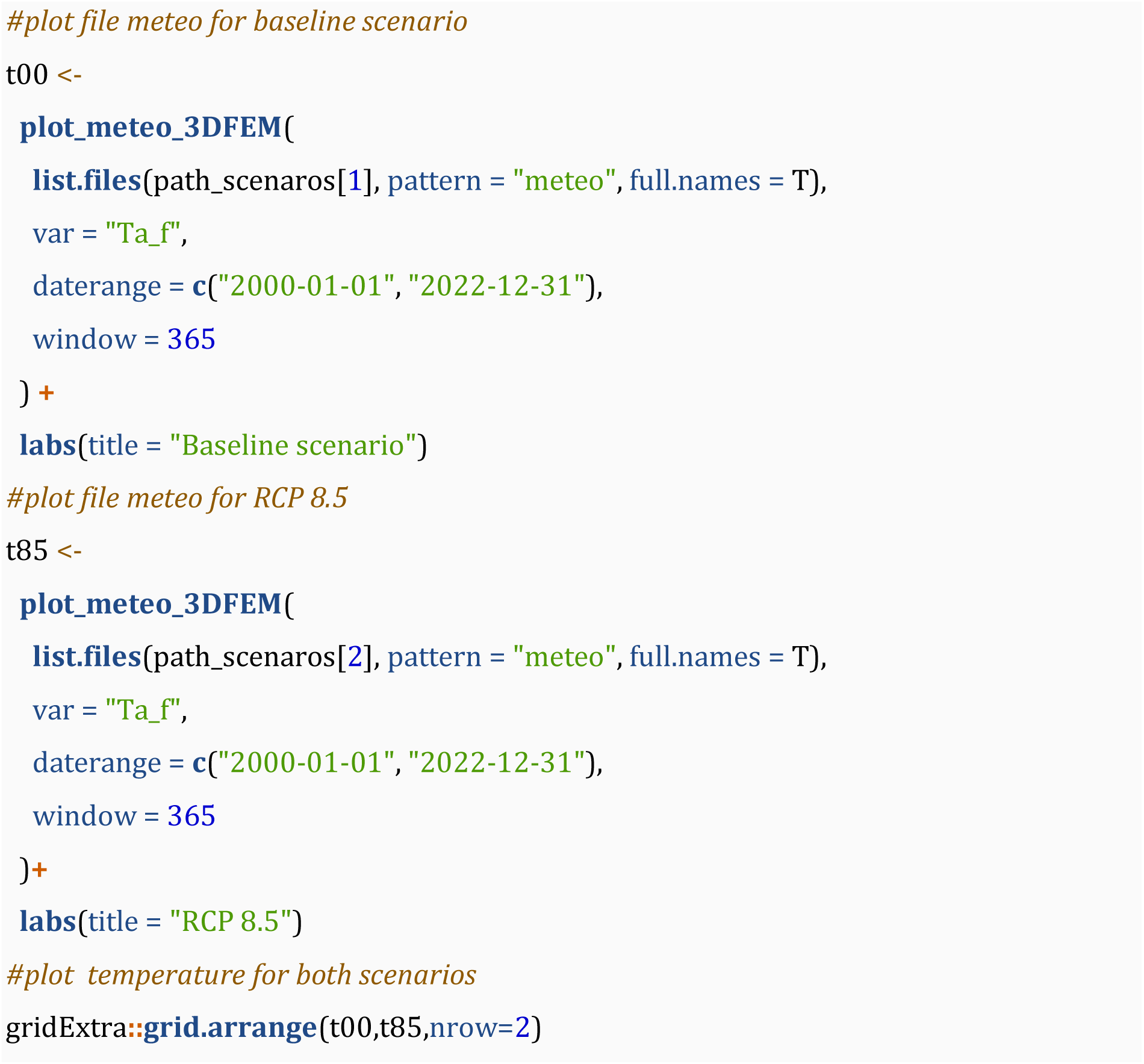

**Figure 5.**
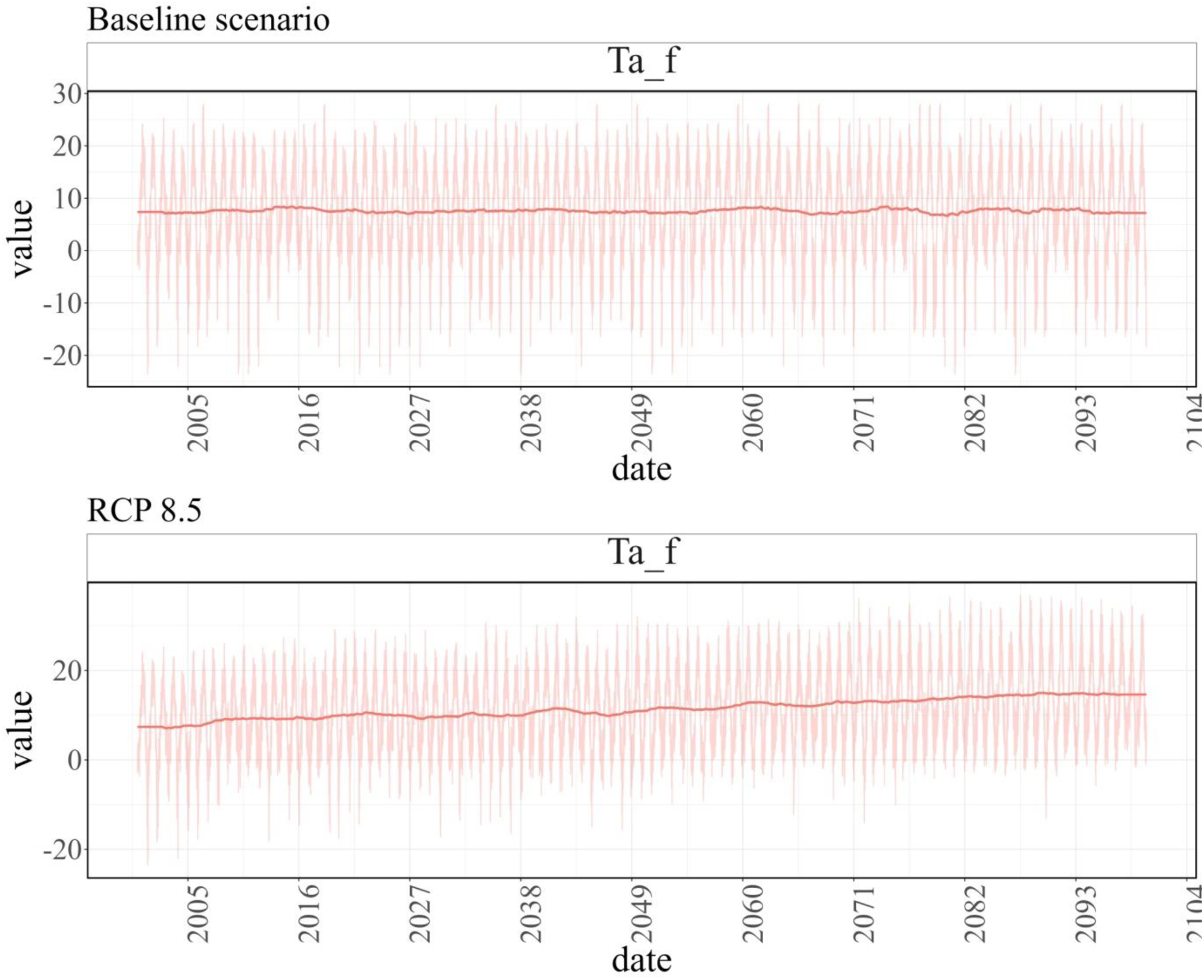
Mean temperature at Bilỳ Křiž under the current climate scenario (baseline, top panel) and the most severe climate change scenario (RCP 8.5, bottom panel).

##### Run simulations

In this piece of code is illustrate a possible way to launch multiple simulations in loop (in this case two simulations). Since each simulation is independent from the others it is possible to launch the simulations in parallel, sending each simulation to a different CPU (not illustrated here).

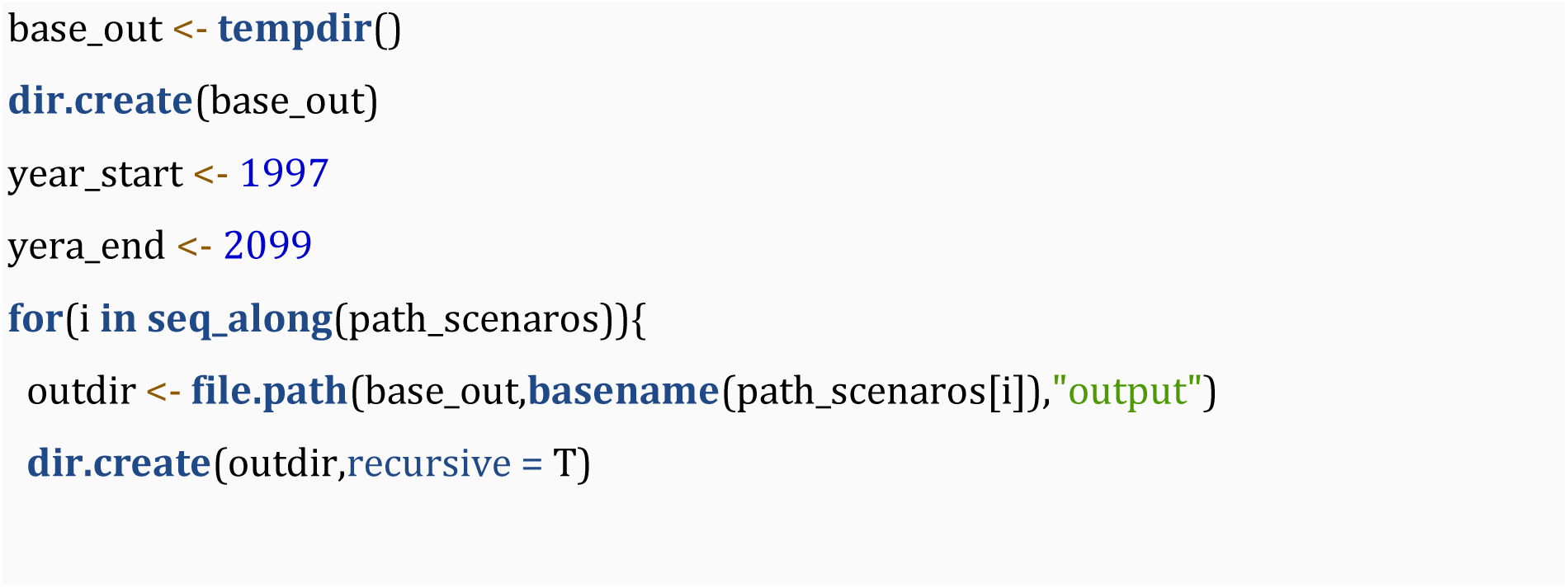
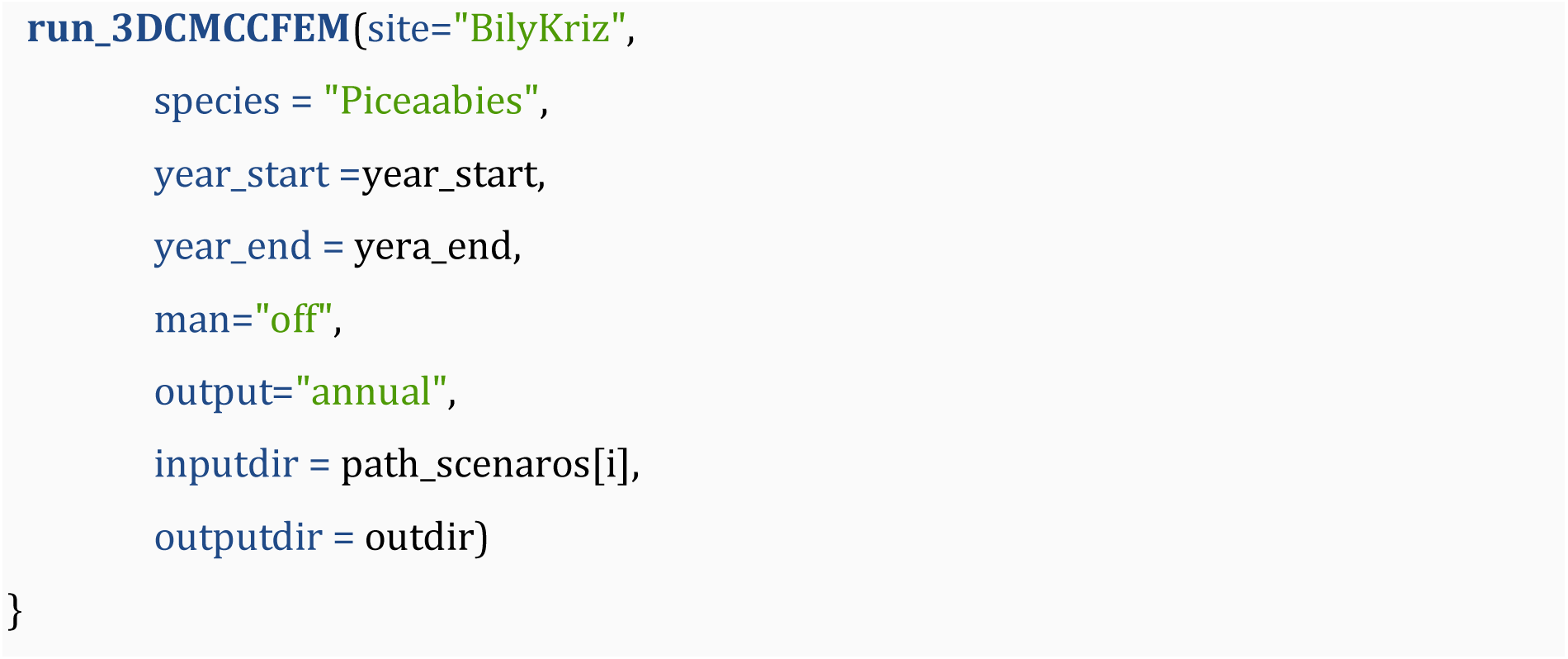

##### Read and plot the outputs

In an analogous way, it is possible to read each output file in loop and to build a single data.frame containing the results of all simulations. In this way, it is possible to compare the different model results in one single plot as shown in Figure 6.

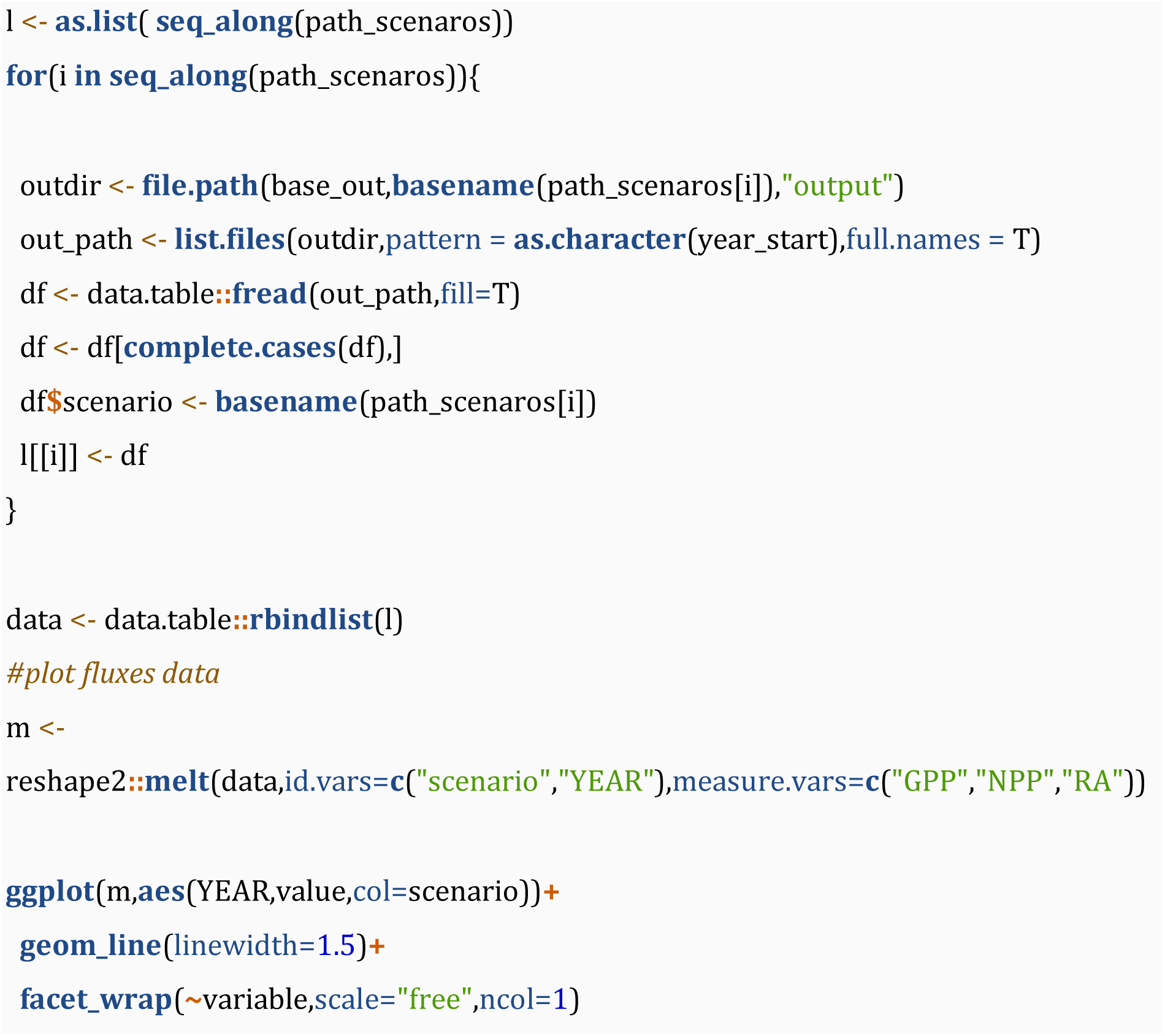

**Figure 6.**
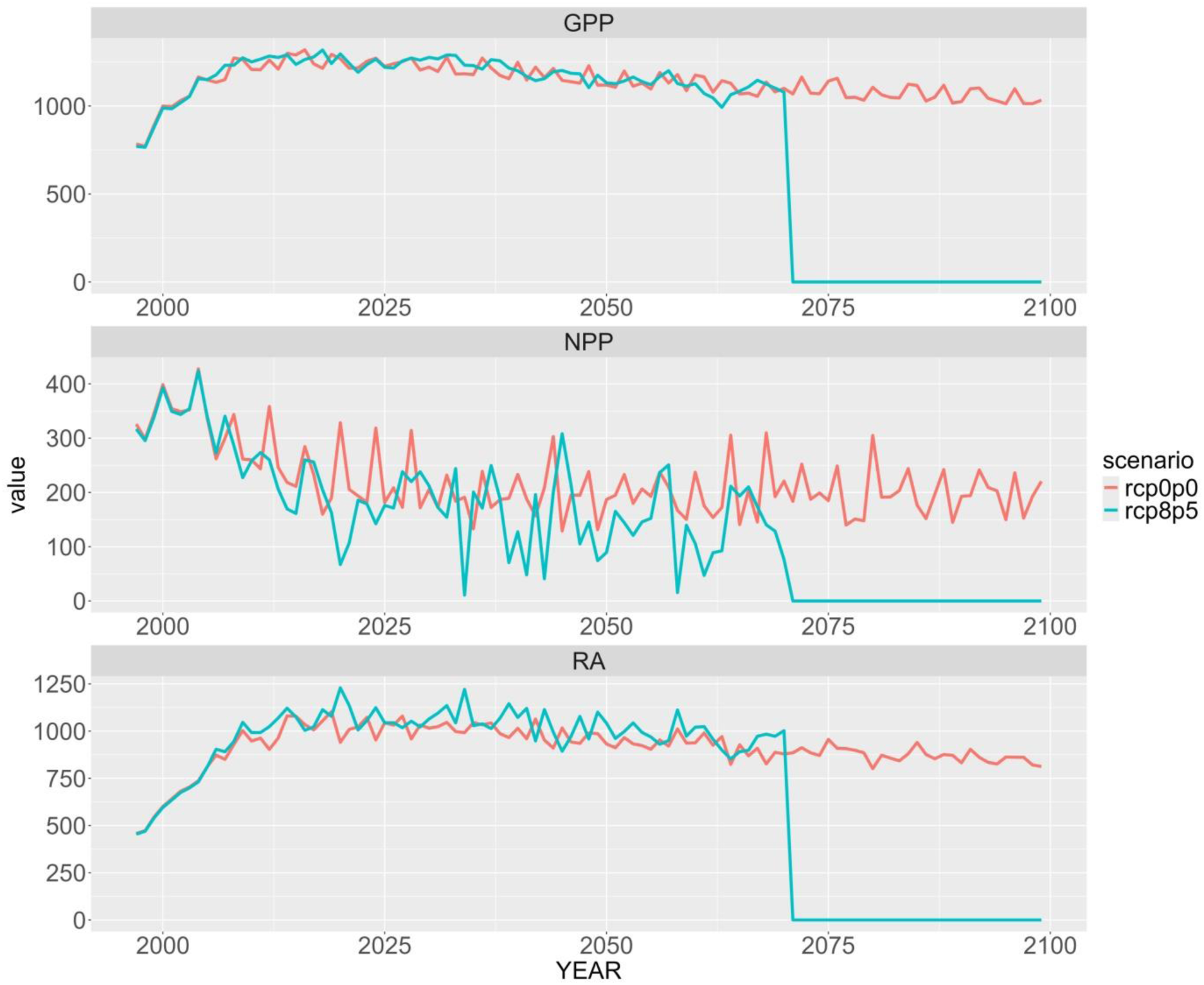
Main fluxes (GPP, NPP, RA; gC m^−2^ day^−1^) at Bilỳ Křiž under the baseline climate and RCP 8.5 scenarios.

In this example under the RCP 8.5 climate scenario, the stand dies because of carbon starvation, i.e. emptying of the carbon reserve pool of the trees. In this case, the variables related to the vegetation incoming fluxes (e.g. photosynthesis) go to 0 while the outgoing fluxes (e.g. heterotrophic respiration due to decomposition) and stocks varying accordingly, and the simulation continues until the expected simulation time frame.

#### 4. Final consideration

PBFMs offer a complementary tool to ground-based forest inventory networks and remote sensing observations for monitoring and predicting future wood and carbon stocks in forest ecosystems and several other variables that are otherwise difficult to measure or monitor continuously. However, the reliability of any model must be verified and tested in different contexts and environments. To do this, as many people as possible should have easier access to these tools. With this aim, we wrap the biogeochemical, biophysical, process-based model 3D-CMCC-FEM in an R-package hoping to expand its use to a wider range of users. Our package provides a ready-to-use tool that allows for quick control of inputs and outputs, in an accessible programming language like R, ultimately simplifying the use of the 3D-CMCC-FEM model, allowing more researchers to make the most from the PBFM capabilities. The simplicity of the developed functions allows even people with minimal knowledge of the R programming language to successfully interact with the model. Despite this advantage we want to stress the necessity to fully understand the model’s underlying characteristics, processes and their interactions, before approaching it and testing it on real case studies. Interested readers can find up-to-date documentation and studies at the model official web page (https://www.forest-modelling-lab.com/the-3d-cmcc-model) and to the model code repository (https://www.github.com/Forest-Modelling-Lab/3D-CMCC-FEM).

The first version of the presented *R3DFEM* package was recently used to investigate the impact of stand age on stability and resilience of forest carbon budget under current and climate change scenarios (Vangi et al., 2024a) and to explore the direct effects of climate change on the total carbon woody stock and mean annual increment across different species and ages cohorts (Vangi et al., 2024b). The 3D-CMCC-FEM model was heavily tested in these years and evaluated for fluxes and stocks in a multitude of forests across Europe, from the plot-level to the regional and national scale and compared against others PBFM (https://www.forest-modelling-lab.com/publications).

## Acknowledgements

The authors want to acknowledge Riccardo Testolin for the first wrap of the model. This research was supported by the following projects: PRIN 2020 (cod 2020E52THS) - Research Projects of National Relevance funded by the Italian Ministry of University and Research entitled: “multi-scale observations to predict Forest response to pollution and climate change” (MULTIFOR, project number 2020E52THS); FORESTNAVIGATOR Horizon Europe research and innovation programme under grant agree-ment No. 101056875; European Union – NextGenerationEU under the National Recovery and Resilience Plan (NRRP), Mission 4 Component 2 Investment 1.4 - Call for tender No. 3138 of December 16, 2021, rectified by Decree n.3175 of December 18, 2021, of the Italian Ministry of University and Research under award Number: Project code CN_00000033, Concession Decree No. 1034 of June 17, 2022 adopted by the Italian Ministry of University and Research, CUP B83C22002930006, Project title “National Biodiversity Future Centre - NBFC”.

## Author’s contributions

All authors have contributed to the package development and drafting of the manuscript. E.V coded the package. All the authors contributed to the interpretation, quality control, and revisions of the package and manuscript.

## Code availability

The source code of the *R3DFEM* package is accessible via GitHub at https://github.com/Forest-Modelling-Lab/R3DFEM. After the download and installation disk occupancy of the *R3DFEM* package is approximately 33MB and it works on Microsoft Windows platforms (Table 2).

**Table 2.**
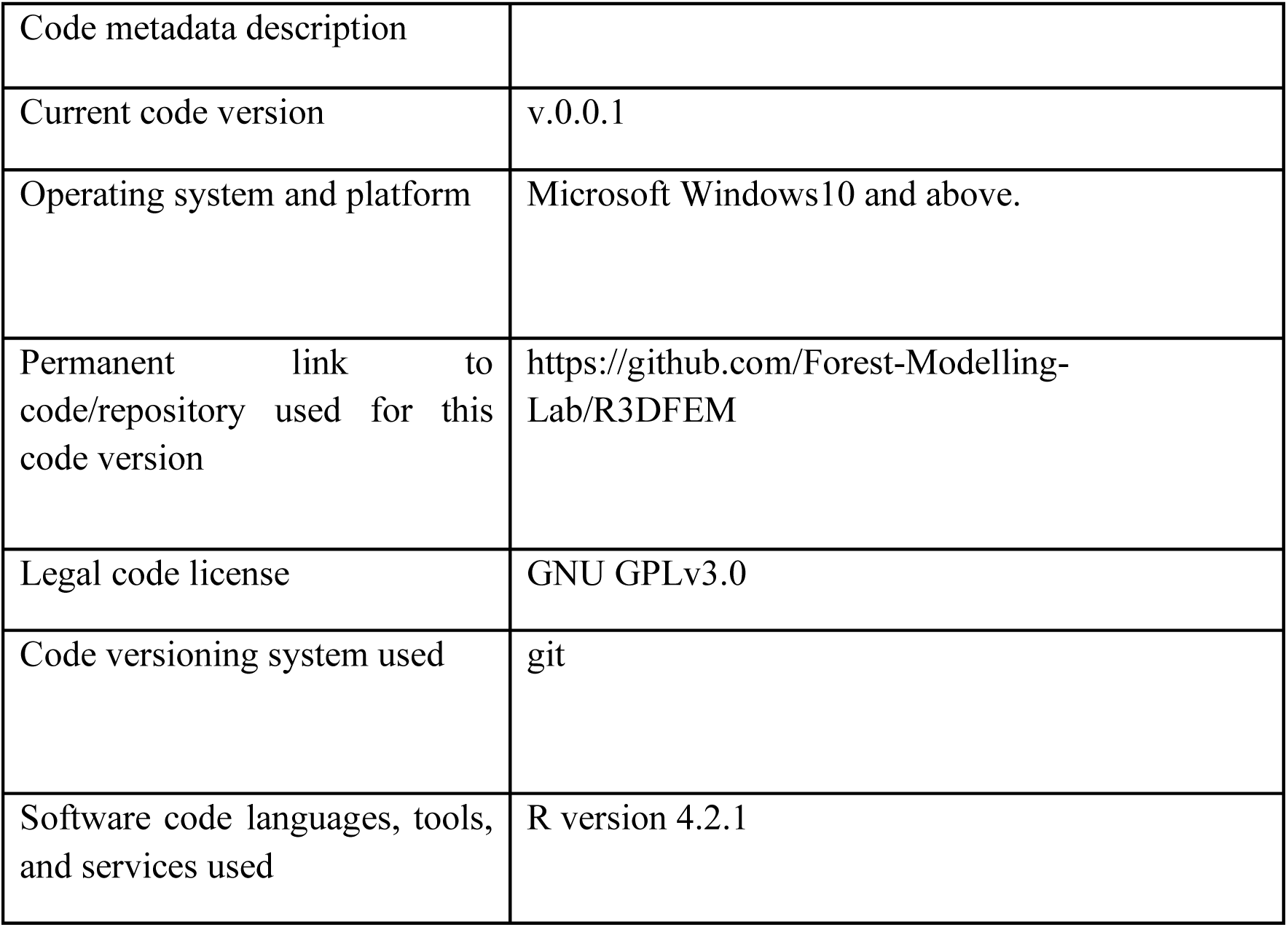
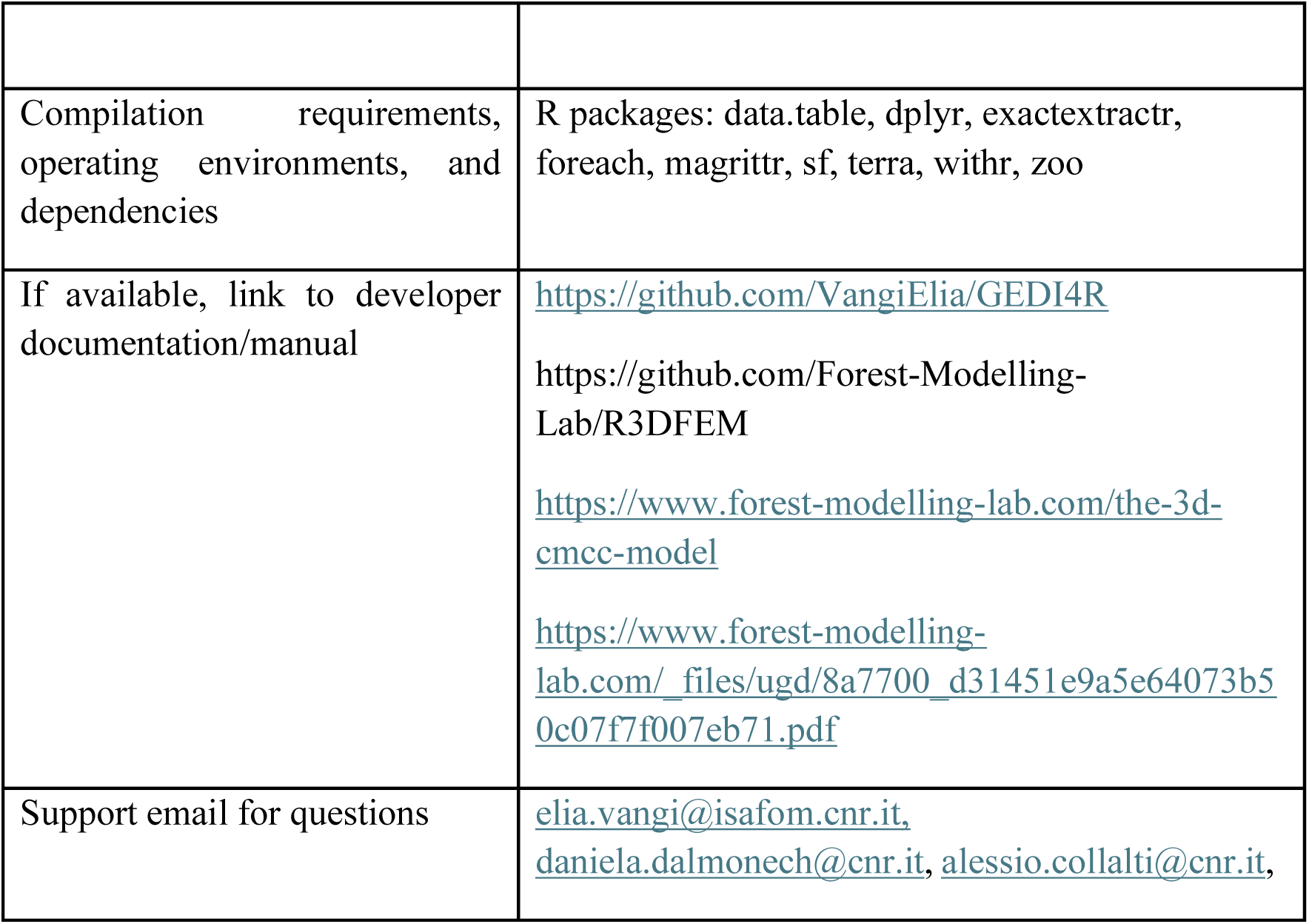
Package metadata

